# Isoform-level transcriptome Atlas of Macrophage Activation

**DOI:** 10.1101/2020.12.20.423532

**Authors:** Apple Cortez Vollmers, Honey E. Mekonen, Sophia Campos, Susan Carpenter, Christopher Vollmers

**Affiliations:** Department of Molecular, Cellular, and Developmental Biology, University of California Santa Cruz, Santa Cruz, California 95064, USA; Department of Biomolecular Engineering, Cellular, and Developmental Biology, University of California Santa Cruz, Santa Cruz, California 95064, USA

## Abstract

RNA-seq is routinely used to measure gene expression changes in response to cell perturbation. Genes that are up or down-regulated following perturbation in RNA-seq studies are designated as target genes for follow-up. However, RNA-seq is limited in its ability to capture the complexity of gene isoforms, defined by the exact composition of exons and transcription start sites (TSS) and poly(A) sites they contain, as well as the expression of these isoforms. Without knowing the composition of the most dominant isoform(s) of a target gene, a minority or non-existent isoform could be selected for follow-up solely based on available annotations for that target gene from databases that are incomplete, or by their nature not tissue specific, or do not provide key information on expression levels. In all, this can lead to loss in valuable resources and time. As the vast majority of genes in the human genome express more than one isoform, there is a great need to identify the complete range of isoforms present for each gene along with their corresponding levels of expression.

Here, using the long-read nanopore-based R2C2 method, we generated an Isoform-level transcriptome Atlas of Macrophage Activation (IAMA) that identifies full-length isoforms in primary human monocyte-derived macrophages (MDMs). Macrophages are critical innate immune cells important for recognizing pathogens through use of Toll-like receptors (TLRs), culminating in the initiation of host defense pathways. We characterized isoforms for most moderate to highly expressed genes in resting and TLR-activated MDMs and generated a user-friendly portal built into the UCSC Genome Browser to explore the data (https://genome.ucsc.edu/s/vollmers/IAMA). Our atlas represents a valuable resource for innate immune research as it provides unprecedented isoform information for primary human macrophages.

## INTRODUCTION

The use of RNA-seq is a primary strategy in biomedical research to identify genes involved in biological processes of interest and how gene expression is impacted upon gene editing or use of chemical or biological agonists. Notably, short-read sequencing technology has reliably been used to quantify changes in gene expression levels or the inclusion level of individual exons and splice junctions. However, because short-read RNA-seq relies on fragmenting RNA molecules prior to sequencing, even advanced computational tools fail at leveraging this ubiquitous data-type into isoform-level information (Bankevich et al., 2012; Grabherr et al., 2011; Pertea et al., 2015). Short-read RNA-seq ultimately falls short in providing comprehensive and accurate full-length isoform structures as well as the level of expression of each isoform under specific conditions.

More recently, long-read technologies from Pacific Biosciences (PacBio) and Oxford Nanopore Technologies (ONT) have made great improvements in sequencing accuracy and throughput and now enable the comprehensive analysis of full-length cDNA molecules at the transcriptome scale (Gupta et al., 2018; Lebrigand et al., 2020; Workman et al., 2019; Wyman et al., n.d.). In contrast to RNA-seq, this technology can determine which isoforms, down to the exact transcription start and poly(A) sites, are expressed at what level by each gene.

The comprehensive transcriptome scale isoform information these technologies provide has the potential to remove the need for targeted and work intensive methods like RT-PCR and 5’/3’ RACE to identify and characterize transcript and/or protein isoforms expressed by a gene. Therefore, comprehensive transcriptome scale isoform information is bound to simplify and improve the outcome of single gene focused follow up studies which include knock-down and knock-out experiments, overexpression assays, Western Blots, ELISAs, pull-downs and many more. This is due to the fact that these assays rely on prior knowledge of what isoform(s) the gene of interest actually expresses in the condition and experimental system being investigated. Finally, detailed knowledge of transcription start sites (TSSs) for each expressed gene in a cell type will also improve the use of CRISPR interference (CRISPRi) technology to knock down genes because guide RNAs can be targeted to TSSs with greater accuracy.

To build on our previous work (Byrne et al., 2017; Robinson et al., 2020) and further push the limits of long-read technology to provide a resource for the innate immune research community, we set out to generate an isoform-level transcriptome atlas of macrophage activation by determining 1) what isoform of a given gene is expressed, 2) at what level, and 3) how isoform and gene expression change following Toll-like Receptor (TLR) activation.

Macrophages are a key cellular component of the innate immune system which represents the first line of host defense against infection and is critical for the development of adaptive immunity (Carpenter et al., 2013; Medzhitov and Horng, 2009). Macrophages recognize conserved structures of microbial-derived molecules or pathogen associated molecular patterns (PAMPs) using TLRs. The regulation of this TLR repertoire fundamentally alters the response to infection (Kawai and Akira, 2008). TLR activation induces the expression of hundreds of genes that encode inflammatory response genes including cytokines, type I interferons, antimicrobial proteins, and regulators of metabolism and regeneration; these molecules in turn mediate inflammation, anti-microbial immunity, and tissue regeneration.

Here we investigated transcriptional responses of human macrophages treated with LPS, Pam3CSK4, R848, and Poly(I:C) which activate TLR4, TLR1/2, TLR7/8, and TLR3, respectively. Using our ONT-based R2C2 method we then generated a total of ~15 million full-length cDNA reads at a median accuracy >99% (Q20) and processed this data into isoforms which we characterized in depth and provide alongside deep Smart-seq2 short-read RNA-seq data as a UCSC Genome Browser session for easy exploration.

## RESULTS

### Experimental setup

To generate a comprehensive isoform-level transcriptome atlas of TLR dependent macrophage activation, we collected PBMCs from two individuals (Rep1 and Rep2) from which we isolated monocytes. From these monocytes, we generated monocyte derived macrophages (MDMs). We treated these MDMs with TLR ligands LPS, Pam3CSK4 (PAM), R848, or poly(I:C) and included a no treatment (NoStim) control. After 6 hours, we collected the stimulated and non-stimulated MDMs and proceeded to extract RNA from each sample. We reverse transcribed the poly(A) fraction of this RNA using a modified oligo(dT) primer and a template switch oligo to generate full-length cDNA with known sequences on both ends. We then amplified this cDNA using PCR and used the resulting double-stranded full-length cDNA as input for both Illumina-based Smart-seq2 (Picelli et al., 2014b) and ONT-based R2C2 (Volden et al., 2018) sequencing protocols (**Fig. 1**).

**Fig. 1:**
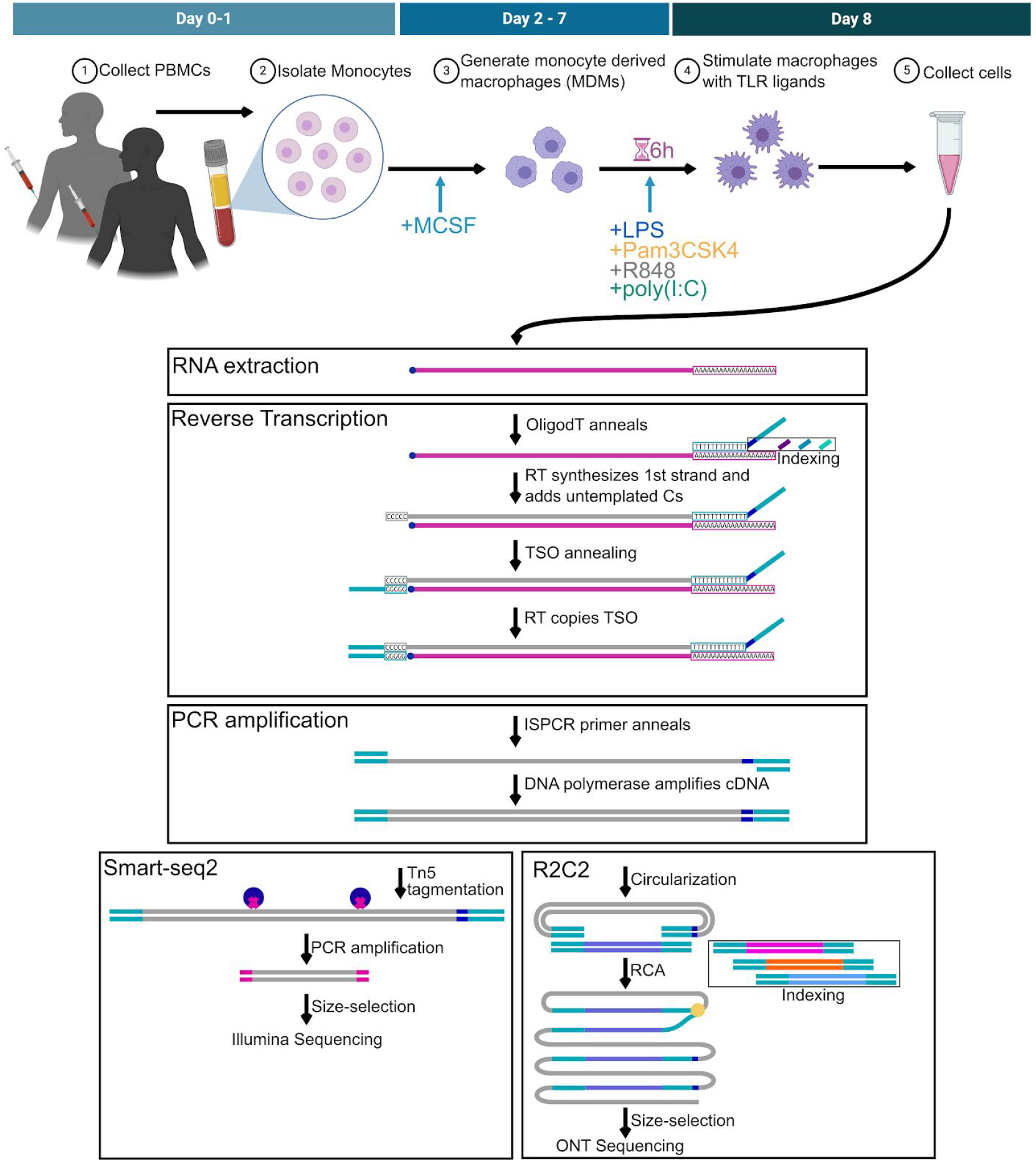
Experimental Design. A schematic of macrophage differentiation and activation is shown on top. A workflow for data generation is shown at the bottom. (Top) We generated monocyte derived macrophages (MDMs) from the peripheral blood mononuclear cells of two individuals (Rep1 and Rep2) by first isolating monocytes and treating them with MCSF. The resulting macrophages were stimulated with TLR ligands for 6 hours and then collected. (Bottom) We extracted RNA and synthesized full-length cDNA which we then processed to generate Smart-seq2 and R2C2 libraries for Illumina and Oxford Nanopore Technologies (ONT) sequencing, respectively. We then performed gene and isoform level analysis of the resulting sequencing data.

### Smart-seq2 based Gene-level Differential Expression Analysis

To identify genes differentially expressed upon TLR activation following treatment with LPS, Pam3CSK4 (PAM), R848, or poly(I:C), we performed Illumina-based Smart-seq2 (Picelli et al., 2014a) sequencing as previously described (Byrne et al., 2019; Cole et al., 2020) (**Fig. S1**, see Methods). We generated approximately 15 - 30 million reads per sample (Table S1), and processed the resulting data using a standard workflow which includes STAR (Dobin et al., 2013), featureCounts (Liao et al., 2014), and DEseq2 (Love et al., 2014). Taking advantage of our biological replicates, we individually compared the LPS, PAM, R848, and poly(I:C) conditions to the NoStim control. Each ligand caused the differential expression of 1000-2000 genes (Table S2 and S3). Between all conditions, 454 genes were shared, a varying number overlapped between three and two stimulants, and some were exclusive to one stimulant (**Fig. 2A**, Table S3). Using Panther Gene Ontology (GO) analysis (Mi et al., 2010; Thomas et al., 2003), we observed that genes shared between all stimulants were strongly enriched for biological processes including “response to cytokine” and inflammatory response” (Table S4). Notably, the genes exclusively responding to R848 were enriched for the “response to organic cyclic compound” biological process, likely because R848 is an organic cyclic compound.

**Figure 2:**
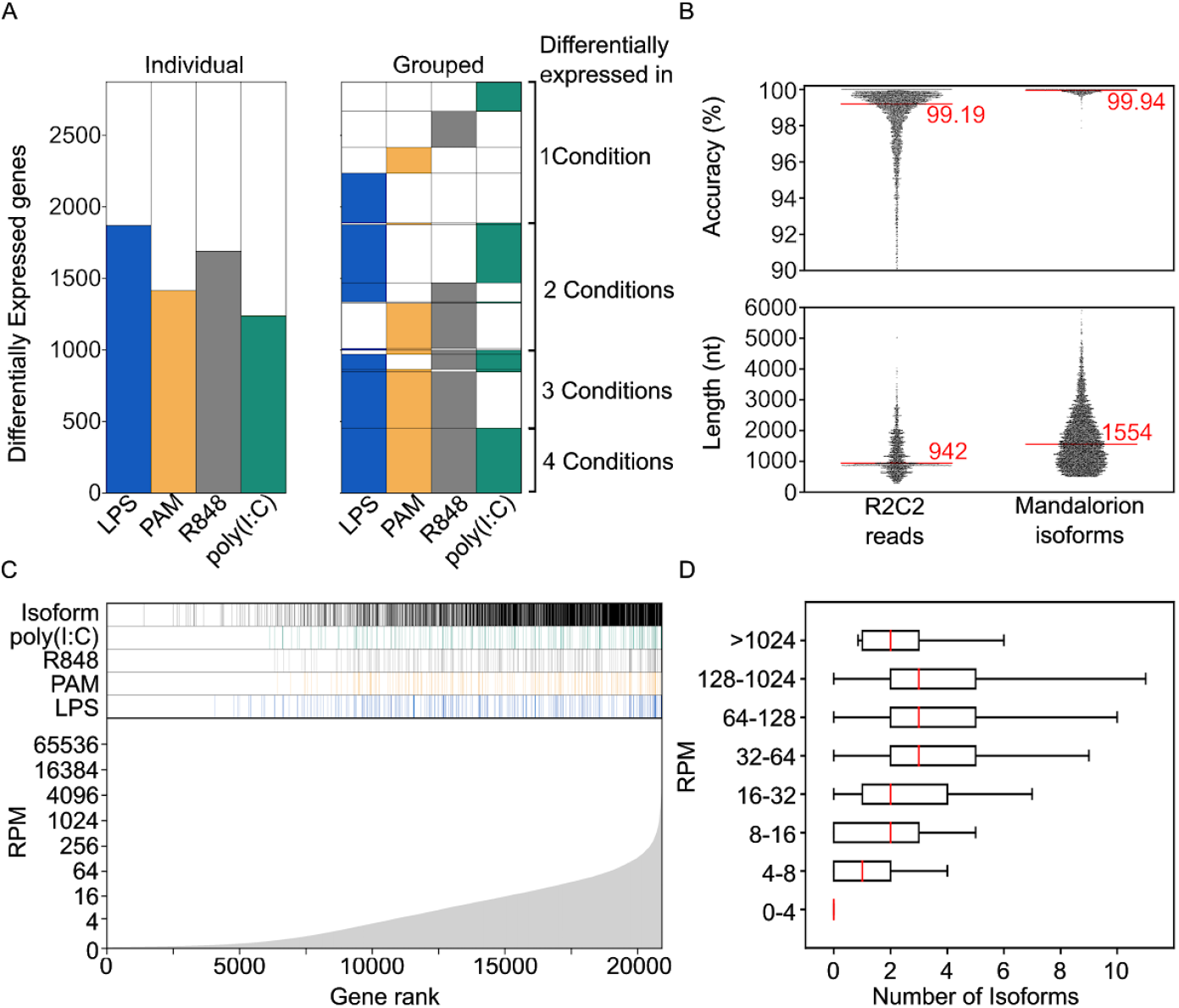
Gene and isoform level analysis. (A) On the left, the numbers of genes differentially expressed between non-stimulated macrophages and macrophages stimulated with the indicated TLR ligands. On the right, genes are grouped if they were differentially expressed in more than one condition. For example, the 454 genes differentially expressed in all four conditions are shown at the bottom. Above those, the number of genes differentially expressed in LPS, PAM, and R848, but not poly(I:C) are shown. (B) Accuracy and length of individual R2C2 reads and Mandalorion isoforms are shown as swarmplots, with median values indicated by red lines and numeric values. (C) Different characteristics of genes ordered by expression are shown in this panel. From the bottom to top this panel shows the average gene expression in reads per million (RPM) across all conditions, whether a gene is differentially expressed following LPS, Pam3CSK4 (PAM), R848, or poly(I:C) stimulation and, whether we identified an isoform for the gene. (D) The number of isoforms we detect for genes is shown as box plots for genes with different expression levels.

### R2C2 based Isoform-level analysis

To supplement this gene-level differential expression analysis, we sequenced the same full-length double-stranded cDNA using our ONT-based R2C2 protocol (Byrne et al., 2019; Volden et al., 2018; Volden and Vollmers, 2020) (**Fig. 1**). R2C2 circularizes cDNA and then amplifies the resulting circular DNA to generate long linear DNA molecules containing concatemeric copies of the initial cDNA. The resulting long DNA is then sequenced and computationally separated into subreads. We then combine these subreads to generate accurate consensus reads of the initial cDNA molecules. To make the creation of this macrophage isoform atlas feasible, we introduced several improvements to the R2C2 method.

First, we enabled multiplexing of cDNA samples by introducing 8nt sample barcodes into the oligo(dT) primers as well as using highly distinct DNA splints for cDNA circularization (**Fig. 1**). Throughout the study, we used these two different indexing strategies to separate technical and biological replicates (Table S5). Indexing samples allowed us to sequence samples in pools on the same ONT flow cells, thereby achieving equal sequencing coverage between samples and minimizing batch effects. Including pilot experiments that sequenced regular and size-selected cDNA of NoStim and LPS samples, we generated 14,961,450 R2C2 reads at a median length of 942nt across multiple ONT MinION flow cells (**Fig. 2B**).

Second, after performing the pilot experiments, we improved R2C2 per read accuracy by increasing the raw read length of R2C2 libraries. We accomplished this by developing a gentle agarose gel extraction protocol that doubled the raw read length of our R2C2 libraries from ~5kb to ~11kb. Combined with a new ONT basecaller, this increased the median per base accuracy of our R2C2 libraries. Previously, this base accuracy was 97.9% (Cole et al., 2020). Here, our most recent sequencing runs show an increased base accuracy of 99.45% (**Fig. S1**). Overall, and including less accurate pilot experiments, the ~15 million reads generated for this study had a median accuracy of 99.19% (Q21).

Third, to take advantage of this improved accuracy and refine the identification of isoforms from the R2C2 reads we generated, we developed a new version of our Mandalorion pipeline (Episode 3.5 - Rogue Isoform) that, amongst several changes, includes improved consensus generation using the Medaka polishing tool and improved handling of isoform ends. When applying this pipeline on the combined ~15 million read data set, we identified 29,637 high confidence isoforms with a median length of 1554nt and a per base accuracy of 99.94% (Q32) which matches the current state-of-the-art for ONT-only consensus accuracy (Shafin et al., 2020) (**Fig. 2B**).

### Enriching DE data set with isoform-level information

Next, we determined which genes these 29,637 isoforms were transcribed from and to what extent isoform identification was dependent on gene expression levels. Our Smart-seq2 based analysis identified 20,915 expressed genes (Average RPM (reads per million) across all conditions and replicates > 0.05). Our R2C2-based analysis identified at least one isoform for 9,688 (46%) of these genes.

Our ability to identify at least one isoform for a gene increased if the gene was expressed at higher levels, i.e. had a higher average RPM (**Fig. 2C**). We identified at least one isoform for 13% of genes between 0.05-3 RPM. This percentage increased with increasing RPM to 49% (3-5 RPM), 64% (5-10 RPM), 80% (10-20 RPM), and 91% (>20 RPM).

Since differentially expressed genes tended to have higher average expression (**Fig. 2C),** we identified at least one isoform for 1986 of the 2873 (69%) genes differential expressed in any condition and 363 of 454 (80%) genes differential expressed in all conditions.

The number of isoforms identified per gene also increased with average RPM (**Fig. 2D**). However, the median number of isoforms per gene did not exceed 3 even in very highly expressed genes. This is likely due to the Mandalorion filtering settings we used which discarded isoforms with less than 1% of all the R2C2 reads at a locus and thereby excluded minority isoforms.

### Identifying genes with differentially expressed isoforms

Next, we established a pipeline to - in addition to gene-level differential expression (DE) - detect genes whose isoforms were differentially expressed between conditions (**Fig. 3A**). While DE pipelines like DEseq2 could be applied to this problem, they would likely just detect isoforms from differentially expressed genes. To detect genes whose relative isoform usage differed between conditions (e.g. Gene A isoform 1 decreases while Gene A isoform 2 increases following stimulation), we applied a Chi-squared contingency table test.

**Fig. 3:**
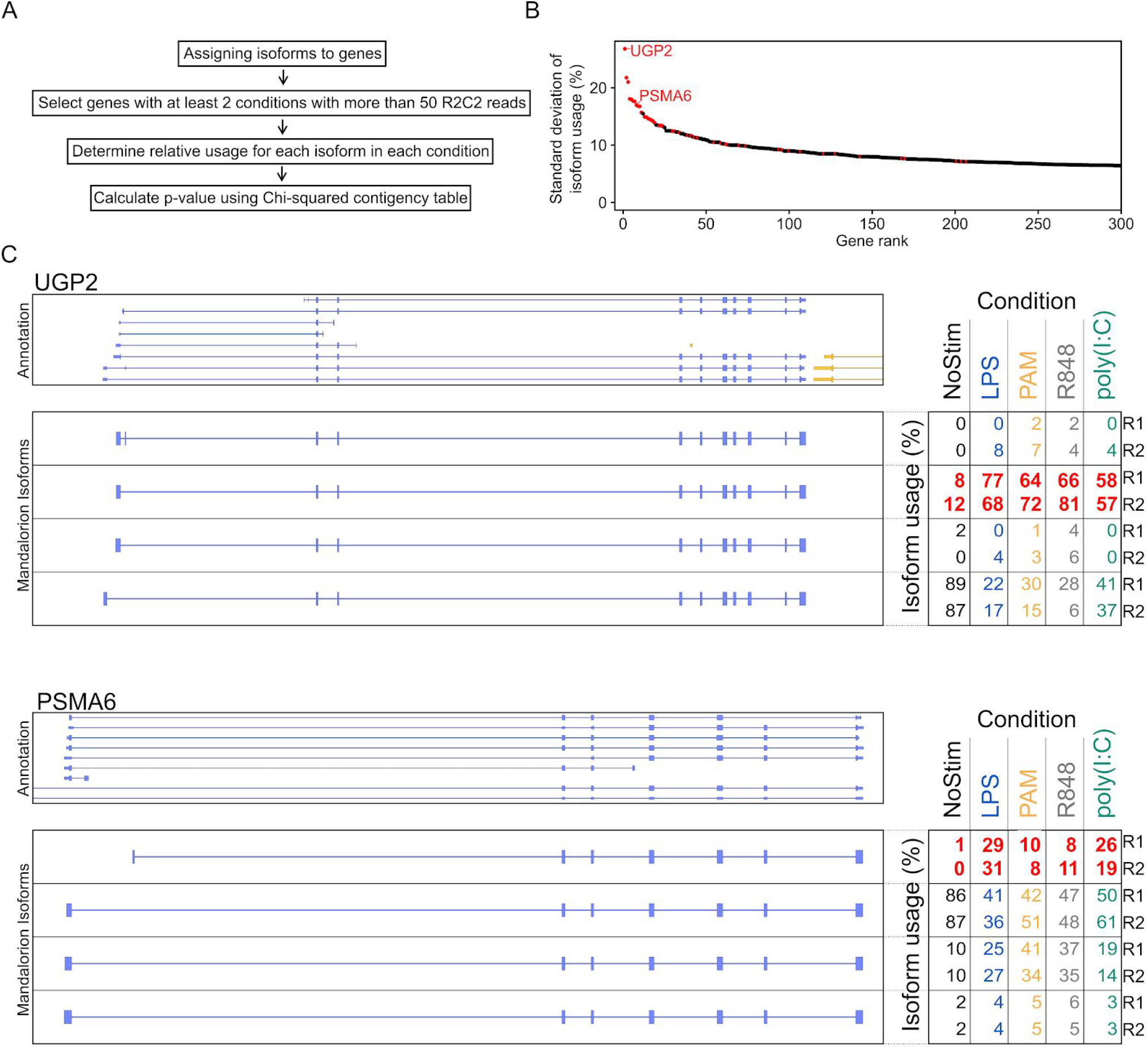
Differential isoform expression. (A) Workflow of differential isoform expression analysis. Genes were sub-selected based on expression and differential expression was determined using a Chi-squared contingency table. (B) Genes are sorted by the maximum standard deviation of relative isoform usage among its isoforms. This maximum standard deviation is plotted (red if the gene has been identified as containing differentially expressed isoforms.) (C) On the left, a Genome Browser view of the indicated genes is shown with GENCODE v34 annotation on top and identified isoforms below. On the right, the relative usage of each isoform in each replicate and condition is shown. Relative usage of the most variable isoform for each gene is highlighted in red.

For each gene, we first determined all the isoforms transcribed from that gene. Then we calculated the relative usage (%) for each isoform in each condition by dividing the number of R2C2 reads associated with that isoform by the number of R2C2 reads associated with all the isoforms in that condition. We then applied the Chi-squared contingency test to the resulting isoform-by-condition relative usage table. To reduce noise, we only tested the 1,872 genes with at least 50 R2C2 reads in at least two experimental conditions. After Bonferroni multiple testing correction, this stringent test produced 47 genes with differential isoform usage with α=0.05 (Table S6). 25 of these 47 (53%) genes were not identified as differentially expressed by the Smart-seq2 short-read workflow.

The 47 genes are likely to contain isoforms whose usage varies strongly between conditions. To confirm this, we determined the standard deviation of this isoform usage between conditions for each isoform in each gene (**Fig. 3B**). By sorting the 1872 genes we tested by the largest standard deviation among their isoforms, we showed that this Chi-square contingency table test did indeed identify genes with isoforms that have highly variable usage between conditions.

The majority of these loci featured alternative transcription start sites (TSSs) which in several cases were located in different and sometimes unannotated first exons (**Fig 3C**, Table S6).

### Classifying isoforms

To evaluate how frequent the use of unannotated exons is across all isoforms and how this affects their coding potential, we first categorized the 29,637 isoforms we identified. To this end, we used the SQANTI (Tardaguila et al., 2018) algorithm which associates isoforms with genes and categorizes them as full-splice matches (FSM), novel in catalog (NIC), novel not in catalog (NNC), incomplete splice-matches (ISM) and other less abundant categories (**Fig. 4A**). FSM and ISM isoforms are defined as fully (FSM) or incompletely (ISM) matching the splice-junction chain of an annotated GENCODE transcript. NIC are defined as isoforms that use annotated splice sites in unannotated configurations. NNCs are defined as isoforms that use at least one unannotated splice site (**Fig. 4A**).

**Fig. 4:**
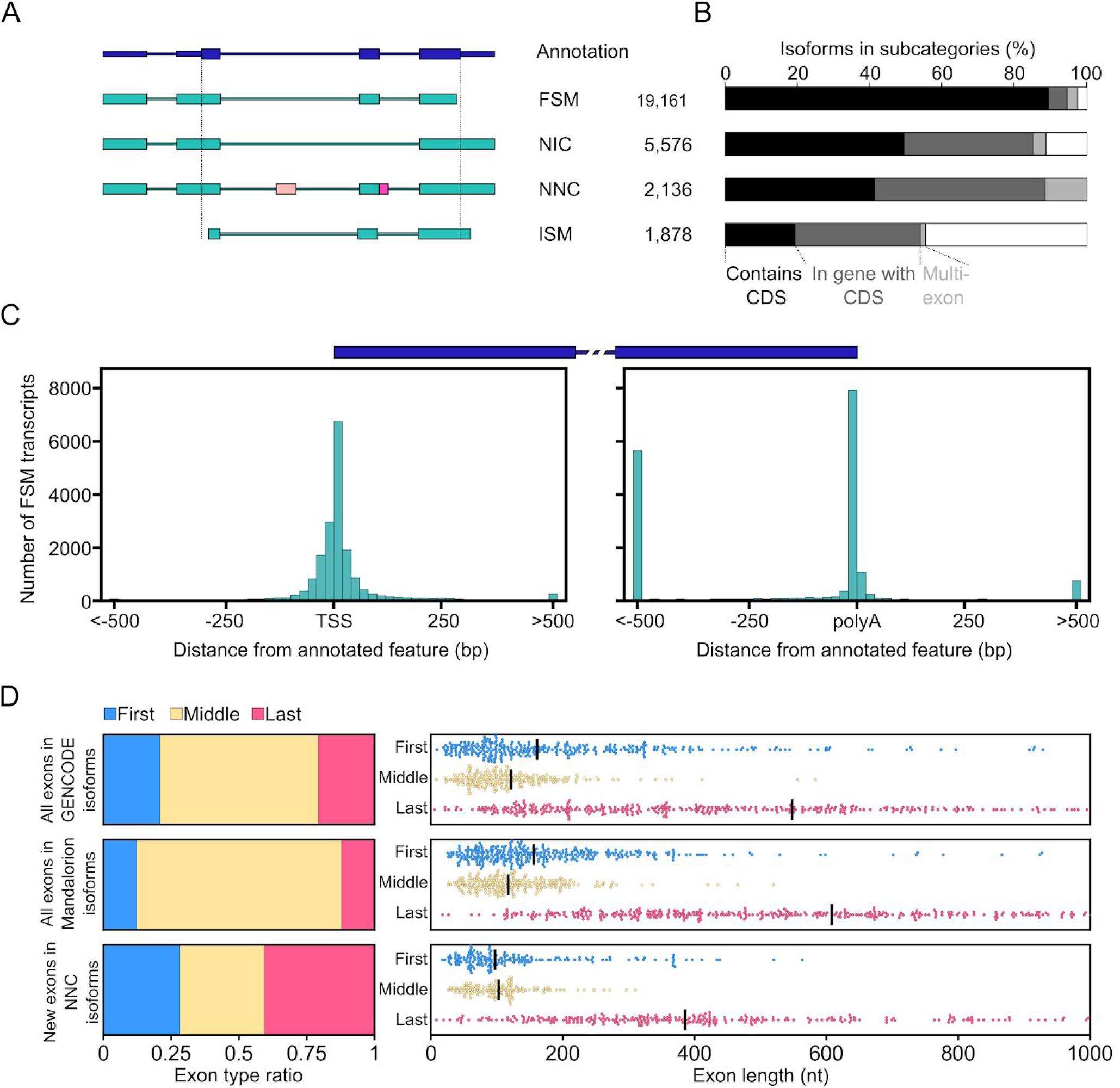
Isoform characterization. (A) On the left, models of different isoform categories are shown. The models shown for Full-splice matches (FSM), novel in catalog (NIC), novel not in catalog (NNC), and incomplete splice-matches (ISM) isoforms all do not contain the CDS of the annotation shown on top. The NNC model contains a new exon (light pink) and an extension of an annotated exon (dark pink). On the right, the numbers of identified isoforms that fall into each category are shown (B) The percentage of isoforms in the different categories that contain more than one exon, fall within a gene that has a CDS, and contain a CDS of that gene are shown as nested bar plots. (C) The distance of 5’ and 3’ ends of FSM isoform to the TSS and TTS of the transcript they are associated with is shown as a histogram. A transcript model is shown on top to give context to the histograms. (D) On the left, the ratio of first, middle, and last exons within GENCODE isoforms, all isoforms identified by Mandalorion, and newly identified exons in NNC isoforms are shown. On the right, the lengths of first, middle, and last exons within these isoform groups are shown as swarm plots with black bars indicating the median.

FSM isoforms represented 65% (19,161), NIC isoforms represented 19% (5,576), NNC isoforms represented 7% (2,136), and ISM isoforms represented 6% (1,878) of the 29,637 isoforms we identified (Table S7). If they were associated with a protein-coding gene, isoforms of different categories had different likelihoods to contain a full coding sequence (CDS) of the gene they were associated with. 94% of FSM, 58% of NIC, 47% of NNC, and 36% of ISM isoforms, which contained more than one exon, contained a full CDS of the protein-coding gene they were transcribed from (**Fig. 4B**).

FSM isoforms which did not contain a full CDS likely had to differ from the GENCODE transcript they matched in their TSS and polyA site positions. Indeed, the 5’ and 3’ ends of FSM isoforms varied in their distance to the annotated TSS and polyA sites of the GENCODE transcript they were associated with (**Fig. 4C**). Taking into account that 5’ and 3’ ends of a FSM isoform may be closer to the TSS or polyA site of another GENCODE transcript in their respective gene, we determined that, of the 19,161 FSM isoforms we identified, 276 had 5’ end and 2068 had a 3’ end more than 500nt away from any annotated TSS or polyA site. The larger number of distant 3’ ends compared to 5’ ends in FSM isoforms could at least in part be explained by last exons being much longer on average than first exons (**Fig. 4D**).

NIC isoforms, which did not contain a full CDS of the gene they were associated with, likely contained a new splice junction between Start and Stop codons which would modify a CDS. NNC isoforms which did not contain a full CDS of the gene they were associated with might differ from the CDS by a few base pairs to encode a slightly different splice-junction or contain entirely new exons (**Fig. 4A**).

### Identifying new exons in NNC isoforms

Next, we focused on NNC isoforms to annotate new exons. We defined a new exon as a part of a transcript whose genomic location does not overlap with a known exon at all (**Fig. 4A**). In the 2,136 NNC isoforms we identified 721 new exons of which 203 were first and 294 were last exons. If new exons were distributed equally among the exons of the 29,637 isoforms we identified, we would expect 89 new first and last exons, indicating that first and last exons are over-represented in this set of new exons (**Fig. 4D**, left).

Further, these new exons were shorter than exons in the GENCODE annotation (v34) or all exons identified by Mandalorion (**Fig 4D**, right). Importantly, the length of new first, middle, and last exons followed the trend of annotated exons with last exons being 2-3x longer than first and middle exons.

Finally, the vast majority of new exons could be validated with short read Smart-seq2 data. Splice junctions leading into these exons (one for first/last exons, two for internal exons) were present in Smart-seq2 reads generated from the same cDNA pool in 665 of 721 (92%) exons. This established that these new exons are highly likely to be present in the cDNA we generated.

### Capturing macrophage-specific lncRNA isoforms

In addition to identifying new exons, NNC isoforms were particularly helpful in redefining long non-coding RNA (lncRNA) loci. 7% of NNC isoforms were associated with lncRNA genes as compared to 3% for both FSM and NIC isoforms. This is likely due to the fact that lncRNA are often expressed at lower levels in a more tissue specific manner than protein-coding genes which complicates their comprehensive annotation (Derrien et al., 2012). Our data set enables the investigation of these lncRNAs and their role in macrophage activation.

In addition to NNC isoforms, isoforms falling into the SQANTI “intergenic” category are also likely to represent non-coding transcripts since protein-coding genes have been exhaustively mapped in the human genome. In total, 184 isoforms were categorized as “intergenic” defined as not overlapping any locus in the Gencode (v34) annotation (Table S7). Of these 184 isoforms, 57 contained more than one exon. These 57 multi-exon isoforms in turn grouped into distinct 38 loci. Only 6 of these 38 loci overlapped with putative lncRNA loci assembled from deep short-read data (Cabili et al., 2011). This showed that investigating specific cell types under different treatments has the potential to identify new previously unobserved lncRNA loci.

### Enabling easy data exploration

While annotations like GENCODE are indispensable for any genomic experiment, they are insufficient when aiming to experimentally follow up potential hits from a screen or an RNA-seq experiment. Our data set addresses this by providing information on which of the potentially numerous isoforms present in (or absent from) the Gencode annotation is actually expressed by a gene of interest and at what level it is expressed compared to other isoforms of that gene. To enable this type of exploration of the data set is available as a custom UCSC genome browser session (https://genome.ucsc.edu/s/vollmers/IAMA). This session contains gene expression information, isoform models and quantification, as well as R2C2 and Smart-seq2 read tracks.

To demonstrate the user interface of the IAMA browser session, we highlight an overlapping pair of lncRNA - LINC01181 and BAALC-AS - which both appear to be differentially upregulated by all TLR ligands based on short-read RNA-seq data. Inspecting these overlapping loci in the genome browser shows that only a very small number of R2C2 reads align to BAALC-AS and that short non-directional RNA-seq reads assigned to BAALC-AS are therefore likely derived from LINC01181 cDNA. Inspecting the isoforms based on these R2C2 reads also shows no match for BAALC-AS but several for LINC01181 (**Fig. 5**). Of the eight spliced isoforms in the LINC01181 locus, none match but some fully contain the annotated LINC01181 transcript. Hovering over the isoforms shows that Isoform_136603_75 is expressed the highest in the LPS condition, taking up 48% of R2C2 reads in that condition. The reference-corrected sequence of Isoform_136603_75 can then be retrieved for down-stream analysis by clicking its model.

**Fig. 5:**
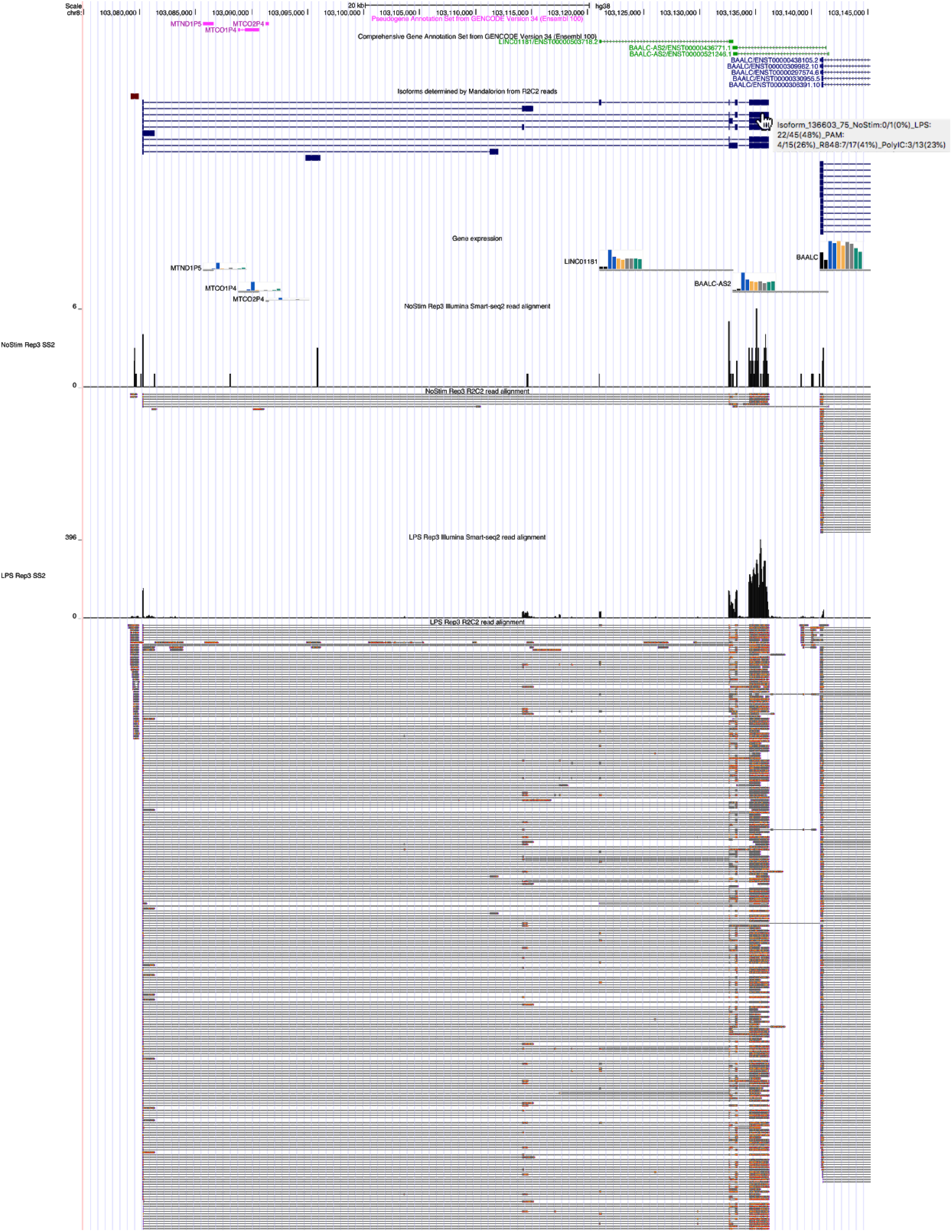
Data exploration. A screenshot of the IAMA session in the UCSC genome browser is shown. From the top, GENCODE annotation, Mandalorion Isoforms, Smart-seq2 based gene expression (bar graphs), Smart-seq2 (histogram), and R2C2 reads. Highlighted are Smart-seq2 and R2C2 reads for just one replicate and two conditions to demonstrate the IAMA browser session and for space-saving purposes. The complete IAMA session for both replicates and all TLR-activated conditions are available here (https://genome.ucsc.edu/s/vollmers/IAMA).

## DISCUSSION

For investigators interested in the in-depth analysis of a gene’s expression and function in a specific cell-type under defined experimental conditions, current efforts to annotate and quantify transcriptomes fall short. This is due to two major limitations of these efforts: 1) the use of short-read RNA-seq methods which precludes the identification and quantification of isoforms, and 2) efforts using long-read methods are mostly limited to the analysis of a limited number of cell-types, almost always at baseline, which will miss isoforms specific to a different cell-type and under different experimental conditions.

The data set and exploration options we present here will be of real-world use to researchers investigating human macrophages at both baseline and after TLR activation and could provide a blueprint for future studies combining short and long-read transcriptome analysis.

The analysis of the transcriptome we generated shows that differential isoform expression between conditions exists but is limited and most often associated with the differential usage of transcription start sites (TSSs) which is similar to observations we have previously made in mouse and human macrophages (Robinson et al., 2020). This shows that the splicing of genes itself is very similar between the conditions we investigated.

We further show that most isoforms we identify match the splice-junction chain of an annotated isoform exactly but often not its TSS and polyA sites. We also detect hundreds of new exons, enriched for first and last exons. The absence of these exons from the Gencode annotation may be caused by technical or biological limitations of previous studies and annotation efforts. Short-read RNA-seq, which is most often used as the foundation for annotation, is characteristically struggling with capturing transcript ends. Further, these new exons might only be expressed in naive or activated macrophages and even then at fairly low levels.

While we believe the entire data set presents a unique window into macrophage biology, the main purpose in creating it was to enable researchers to better understand their gene or genes of interest. We hope that researchers that, for example, might be interested in a particular gene expressed in macrophages after LPS activation will select the isoform with the highest expression in that condition from our data set to synthesize its exact sequence for follow up studies or to, for example, locate its promoter for CRISPRi experiments.

In most cases, the isoform selected from our data set would be different from an isoform picked arbitrarily from an annotation database. Even if the most abundant isoform of a gene was a FSM isoform, it would likely match one of several annotated isoforms for that gene in the GENCODE annotation and might have different TSS and poly sites than the annotated isoform it is matching. Further, in 1028 genes in this data set, the most abundant isoform was a NIC or NNC isoform which by definition aren’t present in the GENCODE annotation. Ultimately, the isoform-level transcriptome we generated here should contain valuable information for the vast majority of medium and highly expressed genes in macrophages.

Finally, while long-read full-length cDNA sequencing is still too expensive to replace routine RNA-seq, the additional isoform-level information provided by full-length cDNA sequencing should make it a valuable addition to target identification and characterization workflows.

## METHODS

### Cell culture

Peripheral blood mononuclear cells (PBMCs) isolated from the buffy coats of healthy blood donors (Stanford Blood Center) were separated by density gradient centrifugation using Ficoll-Paque PLUS (GE Healthcare) followed by 3X washes in HBSS (Sigma Aldrich, H6648), then resuspended in complete RPMI-1640 (Gibco, 11875093) supplemented with 5 mL pen/strep (100X, Gibco, 15140122), 10% FCS (Gibco, 16140-071), 12.5 ml HEPES (1M, Gibco, 15630-080), 5 ml NEAAs (100X, Life Sciences, SH3023801), 5 ml GlutaMax (100X, Gibco, 35050-061), 5 ml Na-Pyruvate (100 mM, Gibco, 11360-070), 500μl ciprofloxacin (10mg/ml, Acros, AC456880050) and plated onto 10-cm tissue culture plated dishes. Non-adherent cells were removed after 2 hours of incubation at 37°C in 5% CO2. The remaining cells were expanded and differentiated into macrophages by culturing cells in the presence of recombinant human M-CSF (R&D, 216-MC-025, 50ng/mL). Cells were cultured for 8 days with the replacement of culture medium every 2 days. Cells were stimulated for 6 h at the following concentrations: LPS (200 ng/ml), Pam3CSK4 (200 ng/ml), poly(I:C) (50 μg/ml), and R848 (1μg/mL). Unstimulated cells were collected at the same time point for use as control.

### Sample Preparations

#### RNA extraction

Total RNA was purified from cells using Direct-zol RNA MiniPrep Kit (Zymo Research, R2072) and TRIzol reagent (Ambion, T9424) according to the manufacturer’s instructions. RNA was assessed for purity using a nanodrop spectrometer (Thermo Fisher). RNA was quantified using a Qubit Fluorometer (Thermo Fisher) and Qubit™ RNA HS Assay Kit (Thermo Fisher, Q32852).

#### cDNA synthesis

For each of these samples, 100-200ng of total RNA was used to generate full-length cDNA using a modified Smart-seq2 protocol(Picelli et al., 2014b). RNA was reverse transcribed using Smartscribe RT (Clontech). For each sample, reverse transcription was primed with a different OligodT primer containing 30 Ts, a 10nt sample index, and a universal ISPCR priming site. The reverse transcription reaction also contained a template switch oligo (TSO-Smart-seq2) to attach the same universal priming site to the 5’ end of transcripts. After reverse transcription, RNA and primer dimers were digested using RNAseA and Lambda Exonuclease (NEB) after which cDNA was amplified using the Kapa Biosystems HiFi HotStart ReadyMix (2X) (KAPA) with the following heat-cycling protocol: 37°C for 30 minutes, 95°C for 30 seconds followed by 12 cycles of (98°C 20 seconds; 67°C 15 seconds; 72°C for 6 minutes). The reaction was then purified using SPRI beads at a 0.65:1 ratio (to retain cDNA longer than 500bp) and eluted in H2O.

#### Smart-seq2

For each of the samples we processed, Smart-seq2 libraries were generated as previously described. In short, full-length cDNA was then tagmented with Tn5 enzyme loaded with Tn5ME-A/R and Tn5ME-B/R adapters. The Tn5 reaction was performed using 50ng of cDNA in 5ul, 1 μl of the loaded Tn5 enzyme, 10 μl of H2O and 4 μl of 5× TAPS-PEG buffer and incubated at 55°C for 5 min. The Tn5 reaction was then inactivated by the addition of 5 μl of 0.2% sodium dodecyl sulphate and 5 μl of the product was then nick-translated at 72°C for 6 min and further amplified using KAPA Hifi Polymerase (KAPA) using a distinct set of indexing primers for each sample and an incubation of 98°C for 30 s, followed by 13 cycles of (98°C for 10 s, 63°C for 30 s, 72°C for 2 min) with a final extension at 72°C for 5 min. The resulting Illumina library was sequenced on an Illumina NextSeq500 1×75 run.

#### R2C2

Amplified cDNA was then sequencing on the Oxford Nanopore Technologies (ONT) MinION sequencer using the R2C2 method(Byrne et al., 2019; Cole et al., 2020; Volden et al., 2018; Volden and Vollmers, 2020). In short, 100ng of cDNA is circularized using 100ng of a DNA splint (Table S8) and 2x NEBuilder HiFi DNA Assembly Master Mix (NEB). This mix was incubated at 50C for 60 minutes. Non-circularized cDNA was digested by adding 5ul of NEBuffer 2, 3ul Exonuclease I, 3ul of Exonuclease III, and 3ul of Lambda Exonuclease (all NEB) and adjusting the volume to 50ul using H2O. This reaction was then incubated 37°C for 16hr followed by a heat inactivation step at 80°C for 20 minutes. Circularized DNA was then extracted using SPRI beads with a size cutoff to eliminate DNA <500 bp (0.65 beads:1 sample) and eluted in 40 μL of ultrapure H2O. Circularized DNA was split into four aliquots of 10 μL, and each aliquot was amplified in its own 50-μL reaction containing Phi29 polymerase (NEB) and exonuclease resistant random hexamers (Thermo) [5 μL of 10× Phi29 Buffer, 2.5 μL of 10 uM (each) dNTPs, 2.5 μL random hexamers (10 uM), 10 μL of DNA, 29 μL ultrapure water, 1 μL of Phi29]. Reactions were incubated at 30 °C overnight. T7 Endonuclease was added directly to each reaction which was then incubated at 37°C for 2h with occasional agitation. The debranched DNA was then size-selected by either using SPRI beads at a 0.5:1 ratio (pilot experiments) or agarose gel extraction.

For the agarose gel extraction, debranched DNA was pooled and concentrated using DNA Clean & Concentrator-5 columns (Zymo Research) and >5kb DNA was then excised from a 1% DNA low-melt agarose gel. Agarose was then melted at 65°C for 10 minutes, transferred to 42°C and digested by immediate addition of 2ul of beta-agaraseI (NEB) per 300ul of melted gel and incubation at 42°C for 1h. Undigested Agarose was then pelleted by centrifugation (14000RPM for 7 minutes in microcentrifuge) and the DNA in the supernatant was extracted using SPRI beads at a 0.7:1 ratio.

The resulting DNA was sequenced on MinION 9.4.1 flowcells. For each run, 1ug of DNA was prepared using the LSK-109 kit according to the manufacturer’s instructions with only minor modifications. End-repair and A-tailing steps were both extended from 5 minutes to 30 minutes. The final ligation step was also extended to 30 minutes. Depending on pore status, runs were DNAseI treated and reloaded after 24 or 48 hours

### Data analysis

#### Smart-seq2

Data in demultiplexed fastq files was aligned to the human genome (hg38) using STAR and standard settings. Gene expression was determined using featureCounts (-O -g gene_name -a) and GENCODE v34(Harrow et al., 2012). Differential gene expression was then determined using DESeq2(Love et al., 2014).

#### R2C2

Data in Fast5 format were basecalled using the bonito research basecaller (version 0.0.5) (https://github.com/nanoporetech/bonito). The resulting fasta files were converted to fastq files by adding a constant Q15 quality score to each base. To generate R2C2 consensus reads we processed and demultiplexed the resulting fastq files using our C3POa pipeline (https://github.com/rvolden/C3POa).

To identify and quantify isoforms we combined data from all samples and used the 3.5 version of Mandalorion (-O 0,40,0,40 -r 0.01 -i 1 -w 1 -n 2 -R 5) (https://github.com/rvolden/Mandalorion) which uses the minimap2 (Li, 2018), racon (Vaser et al., 2017), Medaka (https://github.com/nanoporetech/medaka), and abpoa(Gao et al., 2020) tools. Isoforms were categorized using the sqanti_qc.py script of the SQANTI (Tardaguila et al., 2018). To identify new exons we used the *ProcessSqantiClassification.py* utility of Mandalorion. To investigate protein-coding potential of isoforms, we translated all three possible reading frames of each isoform using BioPython(Cock et al., 2009) and checked whether these translations contained a CDS sequence provided by GENCODE (Harrow et al., 2012). For differential isoform expression we used the Chi-squared contingency test as implemented in SciPy (Virtanen et al., 2020) on data excluding the pilot experiments. For further analysis and visualization of data we used both the Numpy (Harris et al., 2020) and Matplotlib (Hunter, 2007) Python libraries.

## DATA AVAILABILITY

ONT raw sequencing data is available from the Sequence Read Archive (SRA) under Bioproject PRJNA639136. Illumina raw sequencing data is available from the SRA under Bioproject PRJNA660772. Processed data can also be explored as a UCSC Genome Browser session at https://genome.ucsc.edu/s/vollmers/IAMA

## ACKNOWLEDGEMENTS

We thank the Georgia Genomics and Bioinformatics Core (GGBC) for generating and sequencing Smart-seq2 libraries. We acknowledge funding by the National Institute of General Medical Sciences/National Institutes of Health Grant 1R35GM133569 (to C.V.) and R35GM137801 (to Su.C.), NIH Predoctoral Training Grant (T32GM008646), Ford Predoctoral Fellowship, and the Howard Hughes Medical Institute Gilliam Fellowship (to A.C.V.).

## AUTHOR CONTRIBUTIONS

A.C.V. and Su.C. conceptualized the study design. C.V. conceptualized and implemented data generation and analysis strategies. A.C.V., H.E.M., and So.C. performed experiments. A.C.V and C.V analyzed the data. A.C.V, Su.C., and C.V. wrote and edited the manuscript.

---

**Fig. S1:**
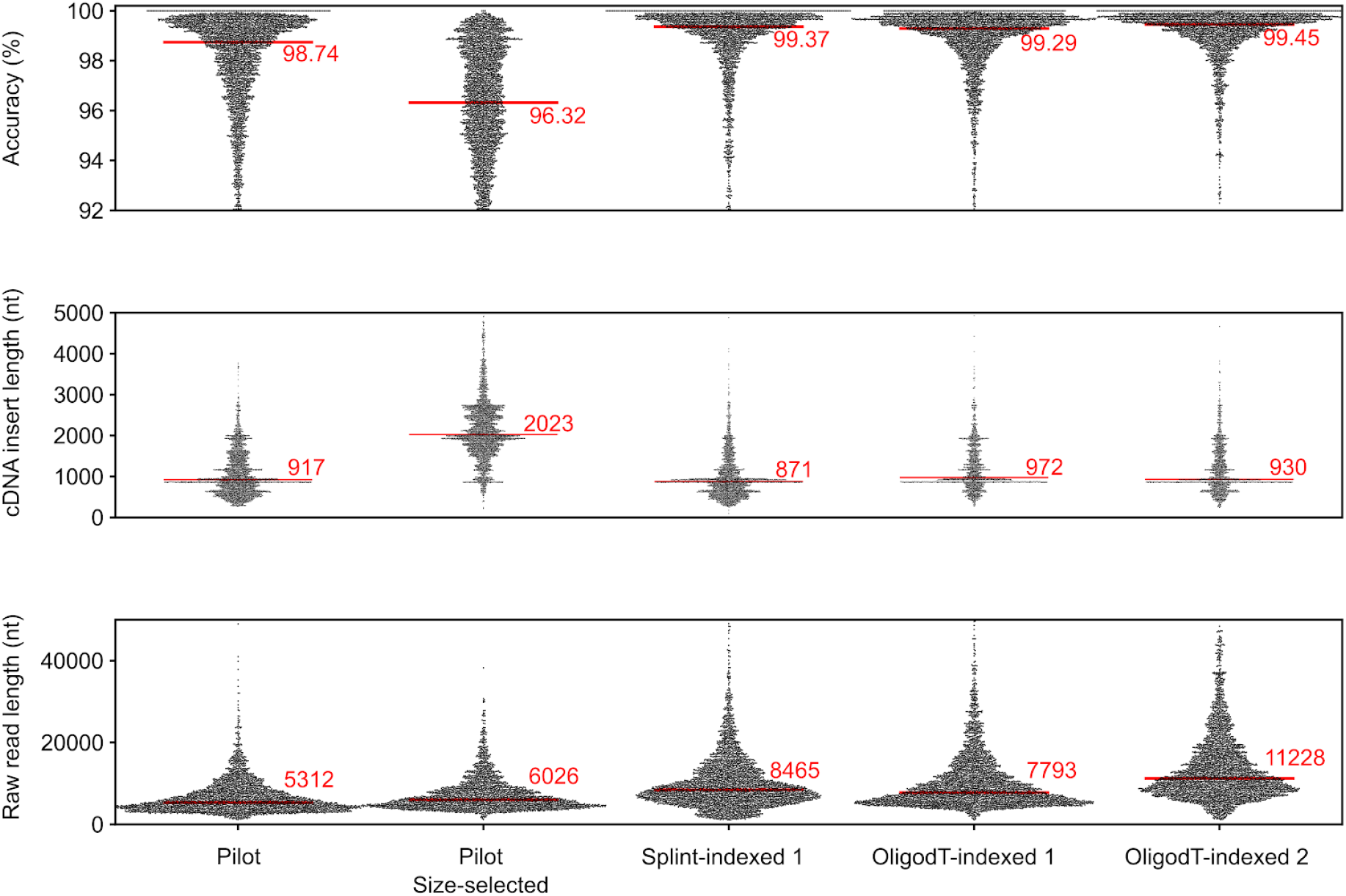
Read characteristics of different R2C2 libraries. Accuracy, cDNA insert length, and ONT raw read length of individual R2C2 reads are shown for different R2C2 libraries as swarmplots, with median values indicated by red lines and numeric values. Splint-indexed 1, OligodT-indexed 1, and OligodT indexed 2 libraries were size-selected by gel excision with OligodT-indexed 2 undergoing the most stringent size-selection.

## REFERENCES

Bankevich A, Nurk S, Antipov D, Gurevich AA, Dvorkin M, Kulikov AS, Lesin VM, Nikolenko SI, Pham S, Prjibelski AD, Pyshkin AV, Sirotkin AV, Vyahhi N, Tesler G, Alekseyev MA, Pevzner PA. 2012. SPAdes: a new genome assembly algorithm and its applications to single-cell sequencing. J Comput Biol 19:455–477.

Byrne A, Beaudin AE, Olsen HE, Jain M, Cole C, Palmer T, DuBois RM, Forsberg EC, Akeson M, Vollmers C. 2017. Nanopore long-read RNAseq reveals widespread transcriptional variation among the surface receptors of individual B cells. Nat Commun 8:16027.

Byrne A, Supple MA, Volden R, Laidre KL, Shapiro B, Vollmers C. 2019. Depletion of Hemoglobin Transcripts and Long-Read Sequencing Improves the Transcriptome Annotation of the Polar Bear (Ursus maritimus). Front Genet 10:643.

Cabili MN, Trapnell C, Goff L, Koziol M, Tazon-Vega B, Regev A, Rinn JL. 2011. Integrative annotation of human large intergenic noncoding RNAs reveals global properties and specific subclasses. Genes Dev 25:1915–1927.

Carpenter S, Aiello D, Atianand MK, Ricci EP, Gandhi P, Hall LL, Byron M, Monks B, Henry-Bezy M, Lawrence JB, O’Neill LAJ, Moore MJ, Caffrey DR, Fitzgerald KA. 2013. A long noncoding RNA mediates both activation and repression of immune response genes. Science 341:789–792.

Cock PJA, Antao T, Chang JT, Chapman BA, Cox CJ, Dalke A, Friedberg I, Hamelryck T, Kauff F, Wilczynski B, de Hoon MJL. 2009. Biopython: freely available Python tools for computational molecular biology and bioinformatics. Bioinformatics 25:1422–1423.

Cole C, Byrne A, Adams M, Volden R, Vollmers C. 2020. Complete characterization of the human immune cell transcriptome using accurate full-length cDNA sequencing. Genome Res 30:589–601.

Derrien T, Johnson R, Bussotti G, Tanzer A, Djebali S, Tilgner H, Guernec G, Martin D, Merkel A, Knowles DG, Lagarde J, Veeravalli L, Ruan X, Ruan Y, Lassmann T, Carninci P, Brown JB, Lipovich L, Gonzalez JM, Thomas M, Davis CA, Shiekhattar R, Gingeras TR, Hubbard TJ, Notredame C, Harrow J, Guigó R. 2012. The GENCODE v7 catalog of human long noncoding RNAs: analysis of their gene structure, evolution, and expression. Genome Res 22:1775–1789.

Dobin A, Davis CA, Schlesinger F, Drenkow J, Zaleski C, Jha S, Batut P, Chaisson M, Gingeras TR. 2013. STAR: ultrafast universal RNA-seq aligner. Bioinformatics 29:15–21.

Gao Y, Liu Y, Ma Y, Liu B, Wang Y, Xing Y. 2020. abPOA: an SIMD-based C library for fast partial order alignment using adaptive band. bioRxiv.

Grabherr MG, Haas BJ, Yassour M, Levin JZ, Thompson DA, Amit I, Adiconis X, Fan L, Raychowdhury R, Zeng Q, Chen Z, Mauceli E, Hacohen N, Gnirke A, Rhind N, di Palma F, Birren BW, Nusbaum C, Lindblad-Toh K, Friedman N, Regev A. 2011. Full-length transcriptome assembly from RNA-Seq data without a reference genome. Nat Biotechnol 29:644–652.

Gupta I, Collier PG, Haase B, Mahfouz A, Joglekar A, Floyd T, Koopmans F, Barres B, Smit AB, Sloan SA, Luo W, Fedrigo O, Ross ME, Tilgner HU. 2018. Single-cell isoform RNA sequencing characterizes isoforms in thousands of cerebellar cells. Nat Biotechnol. doi:10.1038/nbt.4259

Harris CR, Millman KJ, van der Walt SJ, Gommers R, Virtanen P, Cournapeau D, Wieser E, Taylor J, Berg S, Smith NJ, Kern R, Picus M, Hoyer S, van Kerkwijk MH, Brett M, Haldane A, Del Río JF, Wiebe M, Peterson P, Gérard-Marchant P, Sheppard K, Reddy T, Weckesser W, Abbasi H, Gohlke C, Oliphant TE. 2020. Array programming with NumPy. Nature 585:357–362.

Harrow J, Frankish A, Gonzalez JM, Tapanari E, Diekhans M, Kokocinski F, Aken BL, Barrell D, Zadissa A, Searle S, Barnes I, Bignell A, Boychenko V, Hunt T, Kay M, Mukherjee G, Rajan J, Despacio-Reyes G, Saunders G, Steward C, Harte R, Lin M, Howald C, Tanzer A, Derrien T, Chrast J, Walters N, Balasubramanian S, Pei B, Tress M, Rodriguez JM, Ezkurdia I, van Baren J, Brent M, Haussler D, Kellis M, Valencia A, Reymond A, Gerstein M, Guigó R, Hubbard TJ. 2012. GENCODE: the reference human genome annotation for The ENCODE Project. Genome Res 22:1760–1774.

Hunter JD. 2007. Matplotlib: A 2D Graphics Environment. Comput Sci Eng 9:90–95.

Kawai T, Akira S. 2008. Toll-like receptor and RIG-I-like receptor signaling. Ann N Y Acad Sci 1143:1–20.

Lebrigand K, Magnone V, Barbry P, Waldmann R. 2020. High throughput error corrected Nanopore single cell transcriptome sequencing. Nat Commun 11:4025.

Liao Y, Smyth GK, Shi W. 2014. featureCounts: an efficient general purpose program for assigning sequence reads to genomic features. Bioinformatics 30:923–930.

Li H. 2018. Minimap2: pairwise alignment for nucleotide sequences. Bioinformatics 34:3094–3100.

Love MI, Huber W, Anders S. 2014. Moderated estimation of fold change and dispersion for RNA-seq data with DESeq2. Genome Biol 15:550.

Medzhitov R, Horng T. 2009. Transcriptional control of the inflammatory response. Nat Rev Immunol 9:692–703.

Mi H, Dong Q, Muruganujan A, Gaudet P, Lewis S, Thomas PD. 2010. PANTHER version 7: improved phylogenetic trees, orthologs and collaboration with the Gene Ontology Consortium. Nucleic Acids Res 38:D204–10.

Pertea M, Pertea GM, Antonescu CM, Chang T-C, Mendell JT, Salzberg SL. 2015. StringTie enables improved reconstruction of a transcriptome from RNA-seq reads. Nat Biotechnol 33:290–295.

Picelli S, Björklund AK, Reinius B, Sagasser S, Winberg G, Sandberg R. 2014a. Tn5 transposase and tagmentation procedures for massively scaled sequencing projects. Genome Res 24:2033–2040.

Picelli S, Faridani OR, Björklund AK, Winberg G, Sagasser S, Sandberg R. 2014b. Full-length RNA-seq from single cells using Smart-seq2. Nat Protoc 9:171–181.

Robinson EK, Jagannatha P, Covarrubias S, Cattle M, Safavi R, Song R, Viswanathan K, Shapleigh B, Abu-Shumays R, Jain M, Cloonan SM, Wakeland E, Akeson M, Brooks AN, Carpenter S. 2020. Inflammation Drives Alternative First Exon usage to Regulate Immune Genes including a Novel Iron Regulated Isoform of Aim2. bioRxiv. doi:10.1101/2020.07.06.190330

Shafin K, Pesout T, Lorig-Roach R, Haukness M, Olsen HE, Bosworth C, Armstrong J, Tigyi K, Maurer N, Koren S, Sedlazeck FJ, Marschall T, Mayes S, Costa V, Zook JM, Liu KJ, Kilburn D, Sorensen M, Munson KM, Vollger MR, Monlong J, Garrison E, Eichler EE, Salama S, Haussler D, Green RE, Akeson M, Phillippy A, Miga KH, Carnevali P, Jain M, Paten B. 2020. Nanopore sequencing and the Shasta toolkit enable efficient de novo assembly of eleven human genomes. Nat Biotechnol. doi:10.1038/s41587-020-0503-6

Tardaguila M, de la Fuente L, Marti C, Pereira C, Pardo-Palacios FJ, Del Risco H, Ferrell M, Mellado M, Macchietto M, Verheggen K, Edelmann M, Ezkurdia I, Vazquez J, Tress M, Mortazavi A, Martens L, Rodriguez-Navarro S, Moreno-Manzano V, Conesa A. 2018. SQANTI: extensive characterization of long-read transcript sequences for quality control in full-length transcriptome identification and quantification. Genome Res. doi:10.1101/gr.222976.117

Thomas PD, Campbell MJ, Kejariwal A, Mi H, Karlak B, Daverman R, Diemer K, Muruganujan A, Narechania A. 2003. PANTHER: a library of protein families and subfamilies indexed by function. Genome Res 13:2129–2141.

Vaser R, Sović I, Nagarajan N, Šikić M. 2017. Fast and accurate de novo genome assembly from long uncorrected reads. Genome Res 27:737–746.

Virtanen P, Gommers R, Oliphant TE, Haberland M, Reddy T, Cournapeau D, Burovski E, Peterson P, Weckesser W, Bright J, van der Walt SJ, Brett M, Wilson J, Millman KJ, Mayorov N, Nelson ARJ, Jones E, Kern R, Larson E, Carey CJ, Polat İ, Feng Y, Moore EW, VanderPlas J, Laxalde D, Perktold J, Cimrman R, Henriksen I, Quintero EA, Harris CR, Archibald AM, Ribeiro AH, Pedregosa F, van Mulbregt P, SciPy 1.0 Contributors. 2020. SciPy 1.0: fundamental algorithms for scientific computing in Python. Nat Methods 17:261–272.

Volden R, Palmer T, Byrne A, Cole C, Schmitz RJ, Green RE, Vollmers C. 2018. Improving nanopore read accuracy with the R2C2 method enables the sequencing of highly multiplexed full-length single-cell cDNA. Proc Natl Acad Sci U S A. doi:10.1073/pnas.1806447115

Volden R, Vollmers C. 2020. Highly Multiplexed Single-Cell Full-Length cDNA Sequencing of human immune cells with 10X Genomics and R2C2. bioRxiv. doi:10.1101/2020.01.10.902361

Workman RE, Tang AD, Tang PS, Jain M, Tyson JR, Razaghi R, Zuzarte PC, Gilpatrick T, Payne A, Quick J, Sadowski N, Holmes N, de Jesus JG, Jones KL, Soulette CM, Snutch TP, Loman N, Paten B, Loose M, Simpson JT, Olsen HE, Brooks AN, Akeson M, Timp W. 2019. Nanopore native RNA sequencing of a human poly(A) transcriptome. Nat Methods 16:1297–1305.

Wyman D, Balderrama-Gutierrez G, Reese F, Jiang S, Rahmanian S, Zeng W, Others. n.d. A technology-agnostic long-read analysis pipeline for transcriptome discovery and quantification. bioRxiv. 2019.

